# Aird-MSI: a high compression rate and decompression speed format for mass spectrometry imaging data

**DOI:** 10.1101/2025.05.07.652785

**Authors:** Shuochao Li, Hongping Sheng, Pengyuan Du, Jingying Chen, Xixi Wang, Junjie Tong, Jiahua Hong, Xiaohan Jing, Miaoshan Lu, Changbin Yu

**Affiliations:** Central Hospital Affiliated to Shandong First Medical University, Jinan 250000, Shandong Province, China; Alibaba Business School, Hangzhou Normal University, Hangzhou 310000, Zhejiang Province, China; Xi’an Jiaotong-Liverpool University, Suzhou 215000, Jiangsu Province, China

## Abstract

Mass spectrometry imaging has emerged as a pivotal tool in spatial metabolomics, yet its reliance on the imzML format poses critical challenges in data storage, transmission, and computational efficiency. While imzML ensures cross-platform compatibility, its lower compressed binary architecture results in large file sizes and high parsing overhead, hindering cloud-based analysis and real-time visualization.

This study introduces an enhanced Aird compression format optimized for spatial metabolomics through two innovations: (1) a dynamic combinatorial compression algorithm for integer-based encoding of *m/z* and intensity data; (2) a coordinate-separation storage strategy for rapid spatial indexing. Experimental validation on 47 public datasets demonstrated significant performance gains. Compared to imzML, Aird achieved a 70% reduction in storage footprint (mean compression ratio: 30.03%) while maintaining near-lossless data precision (F1-score = 99.26% at 0.1 *ppm m/z* tolerance). For high-precision-controlled datasets, Aird accelerated loading speeds by 15-fold in MZmine.

The Aird format overcomes crucial bottlenecks in spatial metabolomics by harmonizing storage efficiency, computational speed, and analytical precision, reducing I/O latency for large cohorts. By achieving near-native feature detection accuracy, Aird establishes a robust infrastructure for translational applications, including disease biomarker discovery and pharmacokinetic imaging.

## Introduction

Mass Spectrometry Imaging (MSI), a cornerstone technology in spatial metabolomics, has undergone rapid advancement in medical, biological, and environmental sciences owing to its label-free nature and high sensitivity. This technique enables in situ molecular detection with spatial mapping capabilities in biological samples, providing a pivotal tool for resolving spatial heterogeneity in complex metabolic systems.^1–3^ Recent breakthroughs in high-resolution mass spectrometers (e.g., timsTOF flex) coupled with novel ion sources (e.g., AFADESI) have pushed current MSI spatial resolution below 5*µ*m,^4^ whereas nano-secondary ion mass spectrometry (nanoSIMS) achieves sub-100 nm resolution. ^5^ These advancements have driven exponential growth in data dimensionality,^6–8^ with single experiments routinely generating 10^4^-10^5^ pixels. Notably, high-resolution MSI instruments routinely generate hundreds of gigabytes (GB) per large tissue section (more over than 10^5^ pixels),^9^ exacerbating challenges for data storage, transmission and computational analysis. As evidenced by, the METAS-PACE repository, which has accumulated over 12,000 global MSI datasets as of 2024 (Figure 1A), including individual files exceeding 148 GB.^10^ As illustrated in Figure 1B, files exceeding 100 MB constitute over 56% of the total dataset, presenting significant computational challenges for batch processing due to their substantial resource requirements. Innovative data compression algorithms can effectively reduce computational costs and accelerate processing speed in the rapidly developing field of spatial metabolomics.

**Figure 1.**
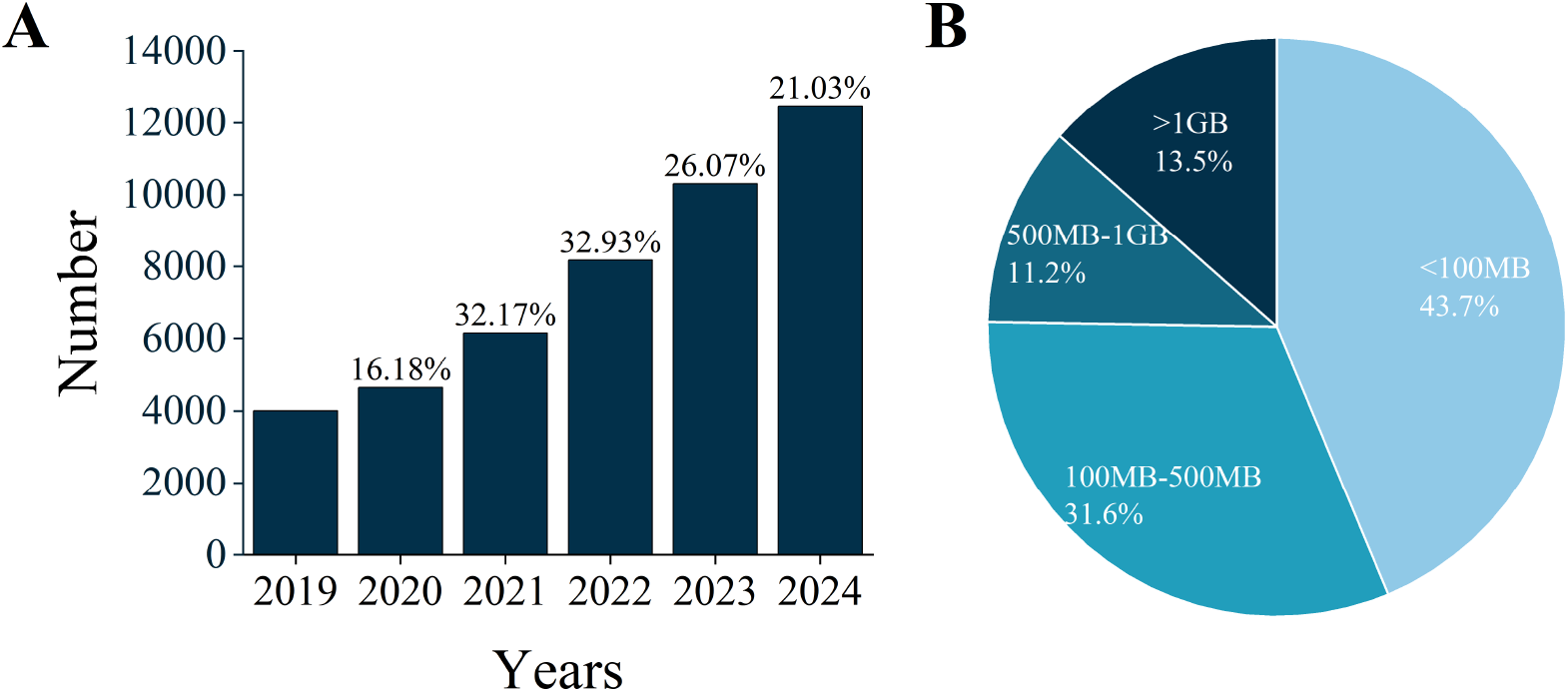
A. The number of files accumulated by METASPACE from 2019 to 2024. The numbers on the bar represent the annual growth rate, the difference of two years divided by the number of last year. B. The situation of the proportion of data in the total data volume after being grouped by size.

The spatial metabolomics data primarily utilize the open imzML(Imaging Mass Spectrometry Markup Language) format,^11^ which implements a dual-file architecture that segregates XML metadata from binary spectral data for cross-platform compatibility. However, imzML exhibits some limitations: (1) lower compressed binary data substantially increases file sizes, thereby elevating network bandwidth consumption during data exchange;(2) high computational overhead during large-file parsing hinders cloud-based analysis and real-time visualization. The lower decoding performance leads to high overhead in decoding large files, which is not conducive to data exchange in cloud computing scenarios and the development of large-scale visualization. Although multiple Mass Spectrometry (MS) data compression algorithms have been proposed,^12–19^ existing solutions failed to simultaneously optimize compression ratios, data precision, and spatial coordinate efficiency, which critically limits their applicability in large-scale spatial metabolomics studies. Aird is an open format for MS data, which features smaller file size and faster read speed. Nevertheless, it doesn’t support MSI data at present.

Herein, we present an enhanced version of the Aird compression format,^17–19^ specifically redesigned for spatial metabolomics through two key innovations: (1) A dynamic combinatorial compression (ComboComp) algorithm for integer encoding of *m/z* and intensity data. (2) A coordinate separation storage strategy enabling rapid spatial indexing. In this study, Aird’s performance in compression ratio, data accuracy, and imaging efficiency was systematically evaluated. The results demonstrated robust capabilities in accelerating mass spectrometry data computation and sharing while maintaining technical reliability for spatial metabolomics workflows. This advancement paves the way for translational applications of spatial metabolomics in critical fields such as disease marker discovery and pharmacokinetics.

## Experimental

### Data Compression and Format

In high-throughput metabolomics, where *m/z* values typically range from 0 to 3000 Da, the Aird format employs an integer transformation strategy to optimize data storage while preserving analytical precision. Based on established precision thresholds (maximum relative error below 2×10^−5^ for *m/z*),^20^ *m/z* values are scaled by 10^5^ and converted to integers, whereas intensity values are accurate to one decimal place or a single digit based on the numerical precision. To mitigate storage inefficiency for non-compressible data segments (e.g., ultra-low-intensity noise signals), raw floating-point formats are preserved. The Aird architecture adopts a dual-file structure comprising: JSON-formatted metadata containing spatially indexed pixel coordinates (X/Y values) extracted from source files (vendor-specific formats or imzML), enabling rapid spatial localization; Mass spectrometry data files compressed through ComboComp, a dynamic combinatorial algorithm that optimizes storage efficiency.

Combocomp advances the concept of combinatorial compression by combining multiple compression algorithms to achieve higher compression rates. The algorithm provides adaptive encoder selection, dynamically allocates algorithm architectures based on numerical distribution characteristics (e.g., successive repetitions, incremental sequences), and supports a variety of algorithms, such as Delta, lz77, and others.^17,18,21^

The AirdPro software suite provides robust conversion capabilities, compressing both vendor-specific raw files and imzML datasets into the Aird format. Key workflow modules are illustrated in Figure 2.

**Figure 2.**
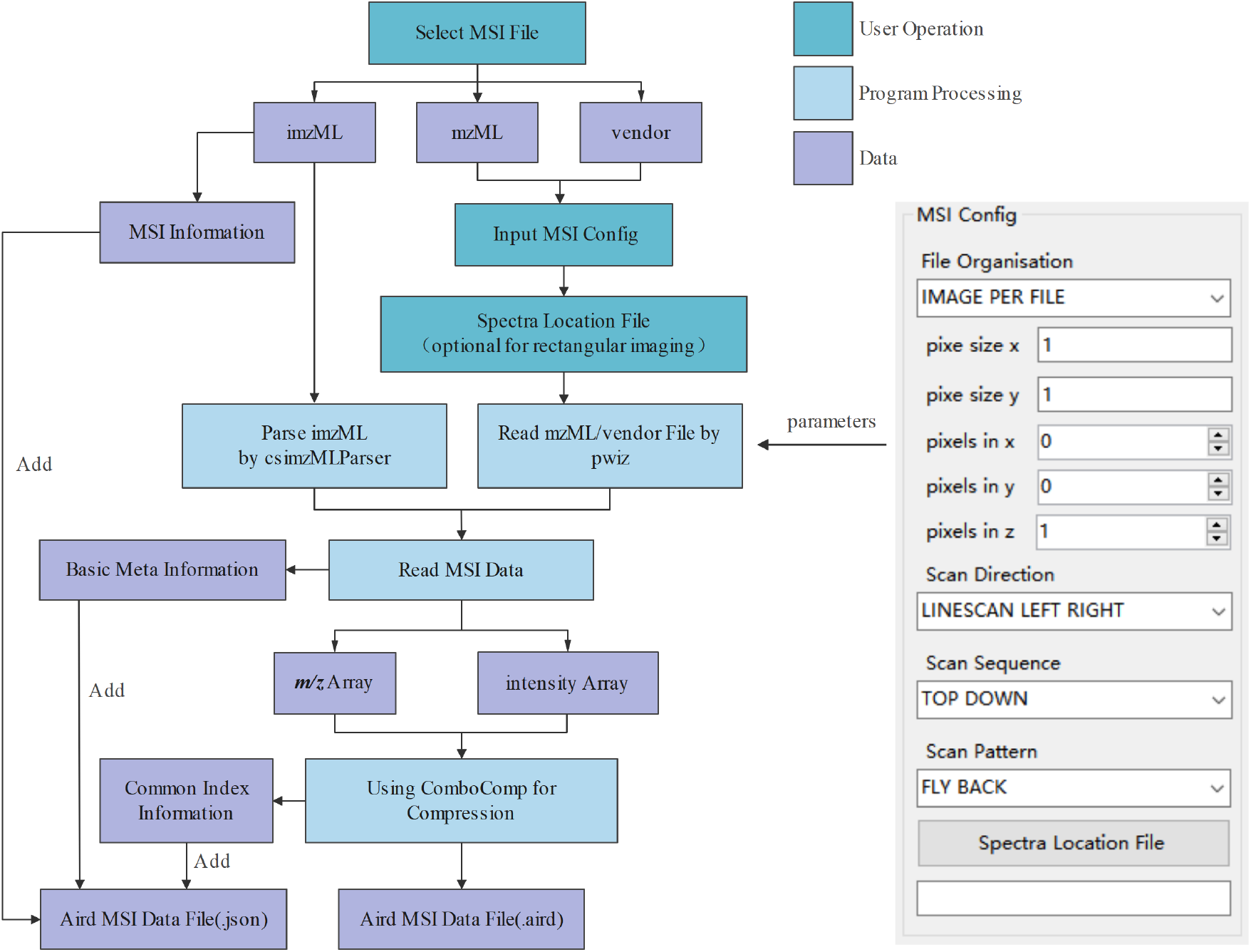
The workflow of AirdPro to convert MSI data, the right side shows the parameters and files that need to be entered when selecting vendor file or mzML file.

### Data Imaging and Visualization

The AirdPro now integrates a spatial metabolomics imaging module that enables real-time visualization of Aird-formatted datasets. The first is efficient decoding, using Aird-SDK to quickly decode coordinate metadata and spectral data, and reconstruct a two-dimensional spatial matrix (dimension: X×Y pixels). This is followed by target *m/z* extraction: userdefined quality queries with adjustable tolerances (*ppm* or absolute error thresholds, such as 3 *ppm* or 0.002 Dalton) to retrieve intensity distributions.

### Experimental Data Preparation

Publicly available mass spectrometry imaging datasets were obtained from two internationally recognized databases: the METASPACE and MetaboLights.^10,22^ The final cohort consisted of 47 datasets (Table S1), which included 12 profile raw files and 35 centroid imzML files, where spatial resolution was 5*µ*m - 100*µ*m, and the file sizes were distributed between 76 MB (MegaByte) and 4 GB (Gigabyte).

All experiments were performed on a dedicated workstation with the following specifications: Windows 10, Intel(R) Xeon(R) Platinum 8260 CPU @ 2.40GHz 2.39 GHz, and 128GB DDR4 memory.

Datasets with Aird file loading times below 1 second in MZmine were excluded from analysis,^23^ as such abbreviated loading durations are subject to software-induced variability that undermines the statistical validity of performance.

### Experimental Design and Performance Evaluation

To validate the compression efficiency and data compatibility of the Aird format, this study systematically evaluated its performance across three dimensions: compression ratio, compression/decompression speed, and data precision. Experimental datasets included original mass spectrometry vendor files (raw format) and imzML files. raw files were initially converted to standard imzML format using imzML Converter (parameters detailed in Table S2), while pre-existing imzML files were compressed into Aird format using AirdPro (parameters detailed in Table S3). During compression, two *m/z* precision modes were implemented: high-precision mode (retaining 5 decimal places [dp], equivalent to 10^−5^) and balanced mode (retaining 4 decimal places, 10^−4^), enabling quantitative assessment of precision loss effects on compression ratios. For all imzML datasets, compression ratios were recorded as:

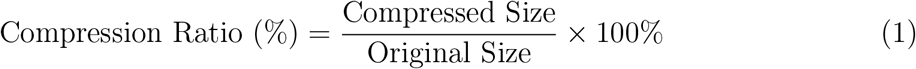

All compression and decompression speed evaluations were conducted on a standardized hardware platform. Compression speed was measured as the time required for format conversion using imzML Converter and AirdPro. Decompression speed was quantified by recording the initial dataset loading time in MZmine. To minimize measurement bias, samples with loading times below 1 second were excluded from analysis.

Feature extraction accuracy was evaluated using MZmine’s MSI feature detection workflow (parameters detailed in Table S4). Metabolite features detected in original imzML files served as the reference dataset. Comparative analysis between Aird and imzML formats was performed using different *m/z* tolerance thresholds (0.1 *ppm*, 5 *ppm*, 10 *ppm*). Consensus features identified within *m/z* tolerance were classified as true positives (TP). Features exclusive to imzML or Aird were designated as false negatives (FN) or false positives (FP), respectively. Performance metrics for each file were calculated as follows:

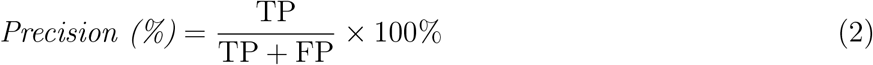

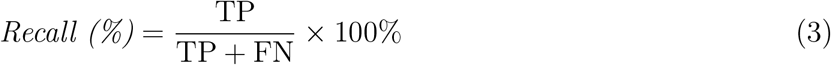

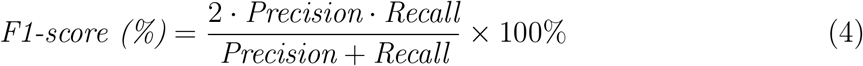

Here, *Precision* (Eq.2) reflects the proportion of correctly identified positive predictions among all positive classifications, *Recall* (Eq.3) quantifies the ability to detect true positives, and *F1-score* (Eq.4) represents the harmonic mean of precision and recall, providing a balanced assessment of classification performance. This multi-threshold approach ensures robust evaluation of format-dependent discrepancies in metabolomic feature detection.

## Results and discussion

### Data Compression Ratios

For raw format files (n=12), compression ratios were evaluated under two precision modes (5dp and 4dp). As shown in Figure 3A, conversion of raw files to imzML format resulted in a 2.9-fold to 3.1-fold increase in file size. In contrast, Aird 5dp achieved a mean compression ratio of 29.98%±0.70%, while Aird 4dp further reduced the mean ratio to 26.98%±0.73% (Figure 3B), demonstrating progressive efficiency gains with precision reduction.

**Figure 3.**
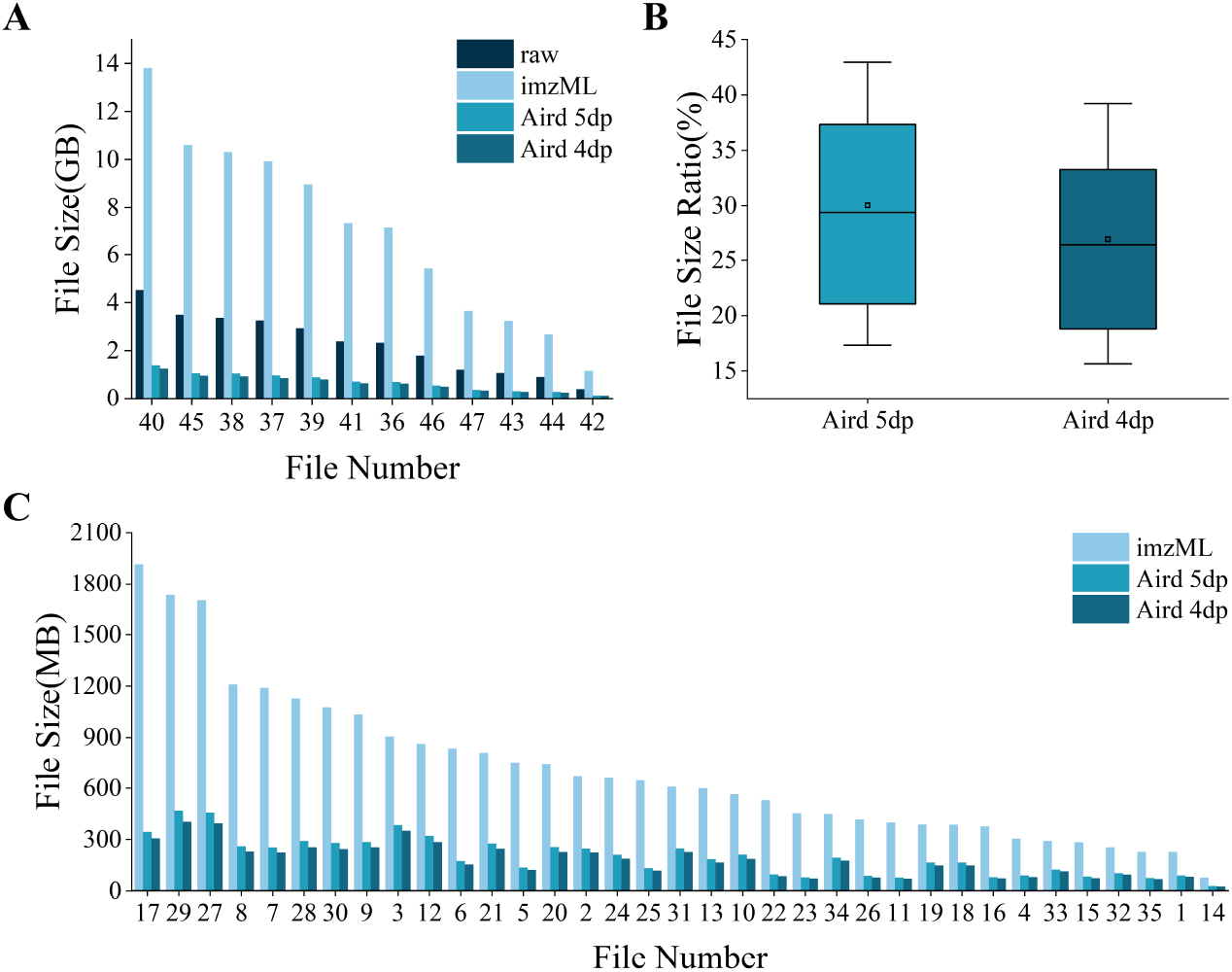
A. Volume comparison of raw format files converted to imzML and Aird formats. B. Compression Ratios (Eq.1) of imzML format files converted to Aird format. C. Volume comparison of imzML format files converted to Aird format.

For imzML format files (n=35), compression performance was tested similarly in both *m/z* precision modes. Statistical analysis revealed significant advantages of the Aird format in spatial metabolomics data compression. Compared to original imzML files, Aird 5dp and 4dp exhibited mean compression ratios of 30.03%±8.73% and 26.89%±8.03% (Figure 3C). The median compressed file size was reduced from 607.68 MB (imzML) to 184.39 MB (Aird), representing a 69.7% reduction in storage and transmission requirements. These results highlight the format’s capacity to balance mass accuracy preservation with substantial storage optimization across diverse instrument platforms.

### Data Compression Time

Conversion times were systematically compared between imzML Converter and AirdPro for transforming raw format data into imzML and Aird formats, respectively. As demonstrated in Figure 4A, the Aird format exhibited significantly faster processing speeds. The mean conversion time for Aird format was 3.27 minutes, representing a 25-fold improvement over the 84.42 minutes required for imzML format conversion. This dramatic efficiency gain underscores Aird’s streamlined encoding architecture and optimized data handling workflows.

**Figure 4.**
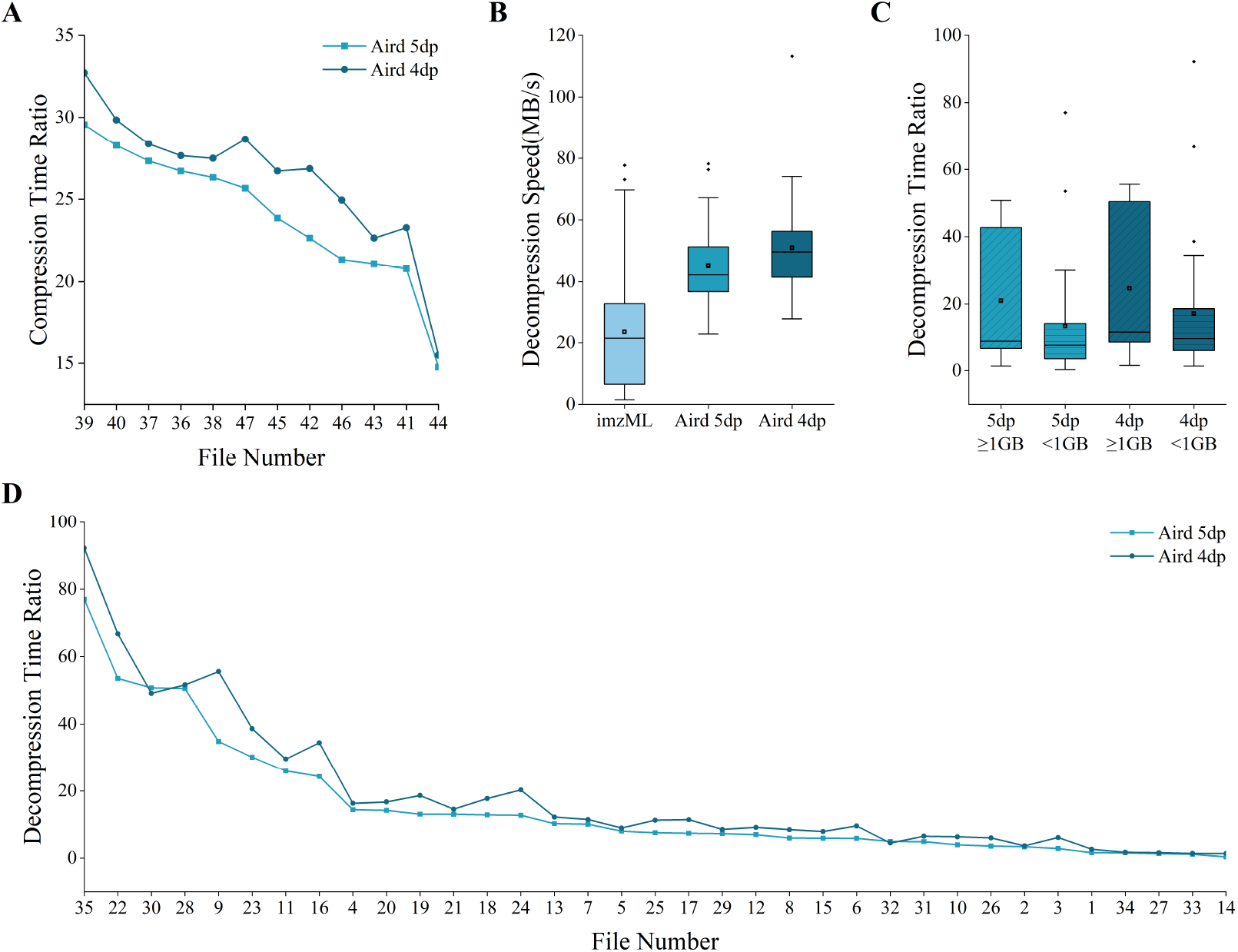
A. Comparison of compression time ratio between Aird 5dp and 4dp. B. Decompression speed in MZmine between Aird and imzML formats. C. Comparison of decompression time ratio stratified by 1GB threshold, showing the significant advantage of Aird in large file scenarios. D. Comparison of decompression time for the two formats in MZmine. The time ratio is calculated by [time(imzML) - time(Aird)]/time(Aird).

### Data Decoding and Loading Performance

File accessibility was rigorously evaluated by comparing loading speeds of Aird and imzML formats in MZmine. Under standardized hardware conditions, the Aird format demonstrated superior decoding efficiency. The mean loading time for Aird 5dp files was 4.37 seconds, achieving a 14.8-fold improvement over imzML’s 68.94 seconds (Figure 4D). While the high-precision mode exhibited about 20% speed reduction compared to the balanced mode, both Aird variants significantly outperformed imzML. As illustrated in Figure 4B, Aird achieved higher data throughput per unit time compared to imzML during file reading operations. Stratifying datasets by a 1 GB file size threshold revealed that Aird exhibited statistically significant improvements in loading efficiency for files exceeding this threshold (Figure 4C). This performance enhancement primarily stems from Aird’s optimized binary encoding architecture, which minimizes redundant data parsing and enables parallelized stream processing.

### Data Accuracy

To assess the precision of Aird’s near-lossless compression algorithm, we systematically compared feature extraction results between Aird and imzML formats using MZmine’s MSI feature detection pipeline.

A systematic comparison of extracted metabolic features across 35 datasets revealed exceptional concordance. At the stringent m/z tolerance threshold of 0.1 *ppm*, the mean F1-score reached 99.26% (Table 1), with individual dataset variations visualized in Figure 5. This near-perfect alignment demonstrates Aird’s high-precision preservation of crucial metabolomic information, achieving over 99% detection concordance with the imzML reference standard. The observed minimal variance in feature intensity distributions further supports Aird’s capability to maintain biochemical relevance while enabling efficient data compression.

**Table 1:**
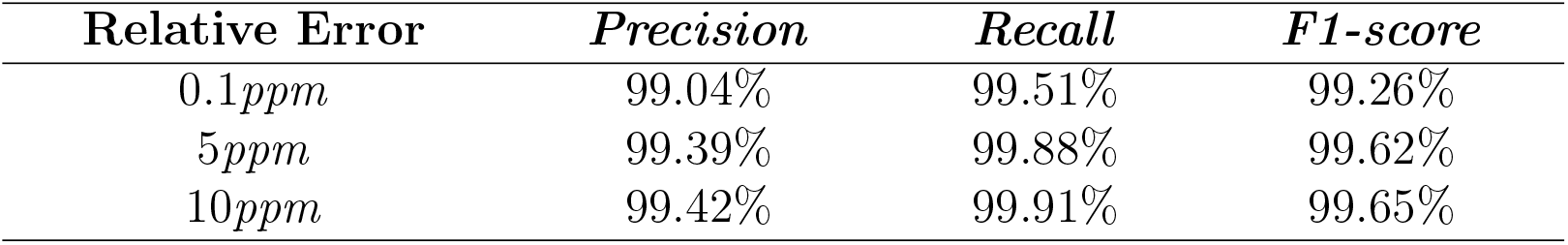
Average *precision, recall*, and *F1-score* of lists on three *m/z* relative error groups processed by MZmine.

| **Relative Error** | ***Precision*** | ***Recall*** | ***F1-score*** |
| --- | --- | --- | --- |
| 0.1*ppm* | 99.04% | 99.51% | 99.26% |
| 5*ppm* | 99.39% | 99.88% | 99.62% |
| 10*ppm* | 99.42% | 99.91% | 99.65% |

**Figure 5.**
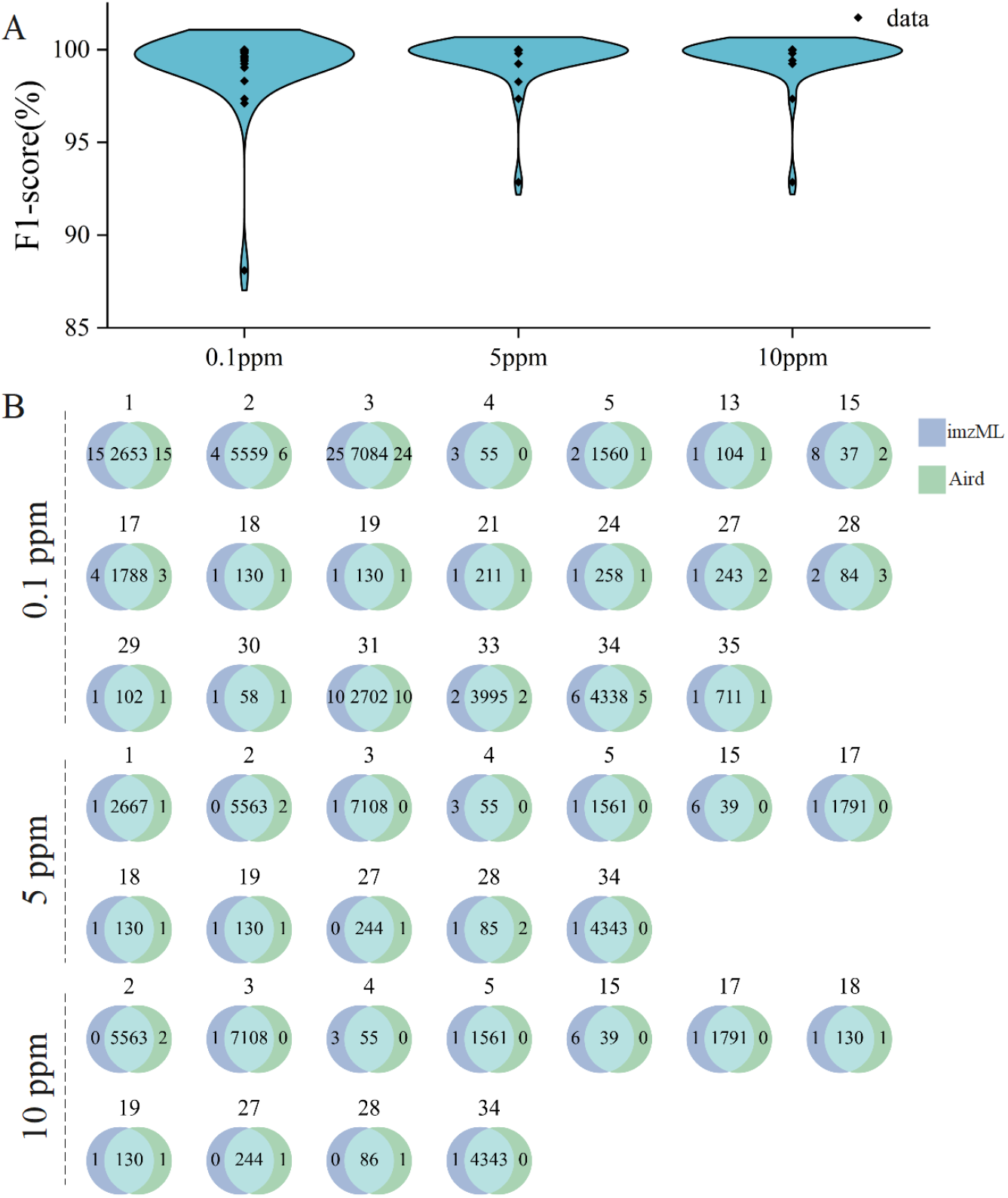
A. F1-score on three relative error groups processed by MZmine feature detection. B. The number of different data features with three *m/z* relative errors.

These results conclusively validate that Aird’s precision-optimized compression architecture introduces no statistically significant deviations in downstream metabolomic analyses, fulfilling the dual requirements of storage efficiency and detection accuracy for spatial metabolomics studies.

### Cross-Platform Imaging Consistency

For a more intuitive comparison, we independently generated the ion images in imzML and Aird formats. The imaging analysis of the dataset S-2210-005030-CMC2pt6_2_S2_MALDI-IF_20241002_IT(Number 17 in Table S1) indicates that The Aird format maintains high visual precision (Figure 6). Among them, the target *m/z* value of 302.0660 (tolerance: 3 *ppm*) was used for linear intensity scaling reconstruction.

**Figure 6.**
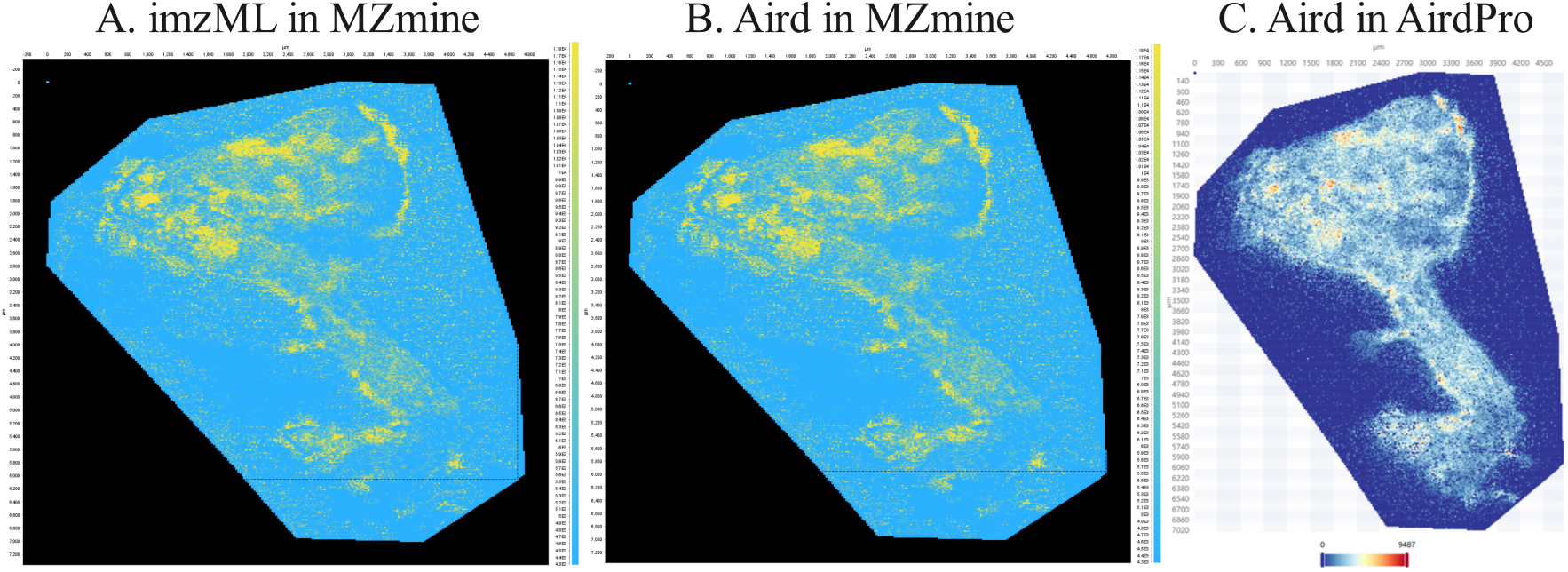
Visual imaging results of the same substance in different software.

### Aird-SDK

The Aird-SDK provides dual functionality for both lossless/lossy compression and high-speed decoding of spatial metabolomics data. This software development kit (SDK) enables customizable processing of mass spectrometry data (including *m/z* values, intensity arrays, and spatial coordinates) under precision-controlled conditions, subject to the following prerequisites: (1)*m/z* : The controlled precision is 10^−5^ with a range less than 2^31^-1, respectively, since most MS data used in metabolomics falls within this range. (2)Intensity: The controlled precision is often 10^0^ with a range less than 2^31^-1.

The Aird format proposed in this study addresses crucial bottlenecks in spatial metabolomics data storage and processing through its ComboComp algorithm and coordinate-decoupled storage architecture. Experimental results demonstrate that Aird achieves a 70% reduction in storage footprint (compression ratio: 30.03% vs. imzML) and a 15-fold acceleration in dataset loading speed, significantly outperforming existing compression frameworks such as mzML (Zlib-based) and mspack.^15^ This performance improvement originates from two synergistic mechanisms: (1) The byte bit count is reduced through integer transformation, where *m/z* and intensity values are converted to integers (*m/z* scaled by 10^5^, intensity by 10^0^), thereby reducing the complexity of floating-point number storage. (2) the ability to read random files, which isolates coordinate metadata (JSON) from spectral binaries to eliminate redundant parsing of spatial headers during pixel-wise access.

Crucially, Aird’s controlled precision loss aligns with established error thresholds for metabolomic feature detection,^20^ ensuring with imzML in downstream analyses. This precision-awareness positions Aird as a viable replacement for imzML in large-scale studies, especially in cases where the bandwidth cost is high during network data exchange.

Aird’s dual focus on file decoding performance beyond simple size reduction. Its 14.8-fold speed advantage over imzML in MZmine loading tasks directly translates to accelerated analytical workflows—a significant improvement for clinical cohorts containing datasets with over 10^6^ pixels.

Future research should prioritize enhancing the ComboComp algorithm’s robustness by integrating advanced spectral decomposition techniques, such as wavelet-based denoising with adaptive thresholding, to mitigate high-frequency noise artifacts while preserving low-abundance ion signatures in mass spectrometry data. Concurrently, the development of AI-driven compression frameworks could address current limitations in static parameterization strategies; specifically, attention-enhanced neural networks trained on tissue-specific metabolic heterogeneity patterns from MALDI-TOF imaging datasets may enable dynamic calibration of *m/z* scaling factors and entropy thresholds, thereby optimizing compression efficiency without compromising spatial resolution. These interdisciplinary advancements would collectively enhance computational reproducibility, scalability, and trustworthiness in large-scale metabolomics workflows.

Current Aird implementations utilize fixed *m/z* scaling factors, which may be suboptimal for ultra-high-resolution instruments. Future implementations incorporating adaptive scaling based on local *m/z* variance could further enhance compression ratios. In summary, Aird redefines spatial metabolomics data infrastructure by balancing rigorous lossy compression with computational efficiency, offering a scalable solution for next-generation biomarker discovery platforms.

## Conclusion

The rapid advancement of spatial metabolomics has driven its widespread adoption in biomedical research, resulting in an urgent demand for efficient data infrastructure to manage exponentially growing datasets. The Aird format addresses this challenge through two synergistic innovations: a dynamic hybrid compression algorithm and coordinate-decoupled storage strategy, achieving 70% storage reduction (average compression ratio: 30.03%) and 15-fold faster decoding speeds compared to imzML. By combining open architecture with adept cloud analytics and real-time visualization, Aird establishes a standardized framework for large-scale data sharing and distributed analysis, positioning it as a transformative tool for accelerating applications in disease biomarker discovery and drug metabolism imaging. Through iterative optimization, the Aird framework is positioned to become the cornerstone infrastructure for spatial metabolomics ecosystems, bridging high-throughput omics technologies with clinical translation to drive paradigm shifts in medicine paradigms.

## Supporting information

Main parameters tables for methods and results in manuscript.

## Acknowledgement

This work is in part of project 2022HWYQ-081, ZR2023LZY009 supported by Shandong Provincial Natural Science Foundation. Academic promotion project of Shandong First Medical University, and funding from Jinan City. The work was supported by the Hainan Province Academician Workstation (Changbin Yu), the specific research fund of The Innovation Platform for Academicians of Hainan Province (No. YSPTZX202407).

## Supporting Information Available

The following files are available free of charge.

- Supplementary.docx: Main parameters tables for methods and results in manuscript.

