## Supplementary material for "Aird-MSI: a high compression rate and decompression speed format for mass spectrometry imaging data": Main parameters tables for methods and results in manuscript.

**Supporting Information Available**

**A. Table**

##### Table S1. MS files used for evaluating the compression rate

| **No** | **Repo** | **Identifier** | **File Name** | **Vendor** | **Format** |
| --- | --- | --- | --- | --- | --- |
| 1 | METASPACE | Agar_w/o_EDC_w_4-APEBA | 20230324_dv_agar_noedc.imzML | Bruker | imzML |
| 2 | METASPACE | Agar_w_EDC_w_4-APEBA | 20230306_dv_agar_paltes_4apeba_all_combined.imzML | Bruker | imzML |
| 3 | METASPACE | Agar_NEDC | 20230307_dv_agar_inter_noapreba_recal_neg.imzML | Bruker | imzML |
| 4 | METASPACE | MPIMM_473_QE_P | 20241010_MS160_slide122_WT638_SDHB_300-1500_20umpixel_A36_pos_402x125.imzML | Thermo | imzML |
| 5 | METASPACE | M28-ALS_4_S4_SM_Neg_20241009_IT | M28-ALS_4_S4_SM_Neg_20241009_IT.imzML | Thermo | imzML |
| 6 | MetaboLights | MTBLS954 | 20160614_DHB_pos_300-2000_100um_154x246_A20_Cacaobean_72h.imzML | Thermo | imzML |
| 7 | MetaboLights | MTBLS954 | 20160913_CHCA_pos_B.puteoserpentis_cryo_350-1400_10um_355x160_A31.imzML | Thermo | imzML |
| 8 | MetaboLights | MTBLS954 | 20160914_DHB_pos_B.puteoserpentis_cryo_350-1400_10um_355x160_A31.imzML | Thermo | imzML |
| 9 | MetaboLights | MTBLS954 | Bathy_puteoserpentis_cryo_DHB_pos_FullMS_475-1200_3um_500x270_A35.imzML | Thermo | imzML |
| 10 | MetaboLights | MTBLS954 | 20161027_DHB_pos_Seagrass_root_FullMS_100-1000_12um_250x150_A30.imzML | Thermo | imzML |
| 11 | MetaboLights | MTBLS954 | 20161125_CHCA_pos_Full450-1200_5um_275x250_A32_MALDI-FISH_B.puteoserpentis7_cryo_run3.imzML | Thermo | imzML |
| 12 | MetaboLights | MTBLS954 | 20181127_MS37_B_child_gill_80_900_DHB_pos_A28_10um_150x390.imzML | Thermo | imzML |
| 13 | MetaboLights | MTBLS954 | 20181219_MS37_B_child_gill_80_900_HCCA_pos_A28_10um_155x420.imzML | Thermo | imzML |
| 14 | MetaboLights | MTBLS954 | 20190123_MS37_MetaboliteMix_Spots_60_900_SDHB_pos_A28_100um_89x74.imzML | Thermo | imzML |
| 15 | MetaboLights | MTBLS954 | BAz3_100x320_10um_E20_1000ms_500-1000.imzML | Thermo | imzML |
| 16 | MetaboLights | MTBLS954 | MS9_20170210_DHB_pos_500-2000_10um_315x200_A28_B.put.imzML | Thermo | imzML |
| 17 | METASPACE | S-2210-005030-CMC2pt6_2_S2_MALDI-IF_20241002_IT | S-2210-005030-CMC2pt6_2_S2_MALDI-IF_20241002_IT.imzML | Thermo | imzML |
| 18 | MetaboLights | MTBLS2639 | earthworm highres.imzML | Bruker | imzML |
| 19 | MetaboLights | MTBLS2639 | section2.3_DHB_100_1000_pos_250x250_A30_8um.imzML | Bruker | imzML |
| 20 | MetaboLights | MTBLS1746 | 20190613_MS1_A19r-18.imzML | Thermo | imzML |
| 21 | MetaboLights | MTBLS1746 | 20190614_MS1_A19r-20.imzML | Thermo | imzML |
| 22 | MetaboLights | MTBLS1746 | 20190822_MS1_A19r-19.imzML | Thermo | imzML |
| 23 | MetaboLights | MTBLS1746 | 20190828_MS1_A19r-22.imzML | Thermo | imzML |
| 24 | MetaboLights | MTBLS1746 | MS1_20180130_PO_1200.imzML | Thermo | imzML |
| 25 | MetaboLights | MTBLS1746 | MS1_20180404_PO_1200.imzML | Thermo | imzML |
| 26 | MetaboLights | MTBLS1746 | MS1_20180405_PO_1200.imzML | Thermo | imzML |
| 27 | METASPACE | 077_biopsy_ITO_timsTOF_1st | 077_ito_timstof_1st.imzML | Bruker | imzML |
| 28 | METASPACE | lung_d119-rll-12b1_nglyc_20240924_bg | lung_d119-rll-12b1_nglyc_20240924_bg.imzML | Bruker | imzML |
| 29 | METASPACE | 077_biopsy_PEN_timsTOF_1st | 077_pen_central.imzML | Bruker | imzML |
| 30 | METASPACE | lung_d227-rll-8c1_nglyc_20240924_bg | lung_d227-rll-8c1_nglyc_20240924_bg.imzML | Bruker | imzML |
| 31 | METASPACE | ecker_slide1_nedc_75um | ecker_slide1_nedc_75um.imzML | Bruker | imzML |
| 32 | METASPACE | T2_tip_c1 | t2_c1_tip.imzML | Bruker | imzML |
| 33 | METASPACE | b5pc_time2_bottom_check | b5pc_time2_bottom_check.imzML | Bruker | imzML |
| 34 | METASPACE | b5pc_time2_top_check2 | b5pc_time2_top_check2.imzML | Bruker | imzML |
| 35 | METASPACE | PNNL05A_V6b_CLMCAFAMM_Lipids_885_1ppm | PNNL05A_V6b_CLMCAFAMM_Lipids_885.imzML | Bruker | imzML |
| 36 | MetaboLights | MTBLS954 | 20160614_DHB_pos_300-2000_100um_154x246_A20_Cacaobean_72h.RAW | Thermo | raw |
| 37 | MetaboLights | MTBLS954 | 20160913_CHCA_pos_B.puteoserpentis_cryo_350-1400_10um_355x160_A31.RAW | Thermo | raw |
| 38 | MetaboLights | MTBLS954 | 20160914_DHB_pos_B.puteoserpentis_cryo_350-1400_10um_355x160_A31.RAW | Thermo | raw |
| 39 | MetaboLights | MTBLS954 | 20161027_DHB_pos_Seagrass_root_FullMS_100-1000_12um_250x150_A30.RAW | Thermo | raw |
| 40 | MetaboLights | MTBLS954 | 20181127_MS37_B_child_gill_80_900_DHB_pos_A28_10um_150x390.RAW | Thermo | raw |
| 41 | MetaboLights | MTBLS954 | 20181219_MS37_B_child_gill_80_900_HCCA_pos_A28_10um_155x420.RAW | Thermo | raw |
| 42 | MetaboLights | MTBLS954 | 20190123_MS37_MetaboliteMix_Spots_60_900_SDHB_pos_A28_100um_89x74.RAW | Thermo | raw |
| 43 | MetaboLights | MTBLS954 | BAz3_100x320_10um_E20_1000ms_500-1000.RAW | Thermo | raw |
| 44 | MetaboLights | MTBLS954 | MS9_20170210_DHB_pos_500-2000_10um_315x200_A28_B.put.RAW | Thermo | raw |
| 45 | MetaboLights | MTBLS1746 | 20190613_MS1_A19r-18.RAW | Thermo | raw |
| 46 | MetaboLights | MTBLS1746 | MS1_20180404_PO_1200.RAW | Thermo | raw |
| 47 | MetaboLights | MTBLS1746 | MS1_20180405_PO_1200.RAW | Thermo | raw |

##### Table S2. The key parameters used for raw data converting in imzML converter(v1.1.4.5i beta)

| **Module Name** | **Key Parameters** | **Contents** |
| --- | --- | --- |
| Imaging | Binary mode | Processed |
|  | Intensity type | Single 32 Bits |
|  | m/z type | Double 64 Bits |
|  | Dimensions | Same as imzML |

##### Table S3. The key parameters used for data converting in AirdPro(v6.0.0.0)

| **Key Parameters** | **Contents** | **Files** |
| --- | --- | --- |
| m/z precision | 5(5dp),4(4dp) | All files |
| Intensity Integer-purpose | VB | All files |
| Intensity General-purpose | ZSTD | All files |
| m/z Integer-purpose | IVB | All files |
| m/z General-purpose | ZSTD | All files |
| Dimensions | Same as imzML | Raw files |

##### Table S4. The key parameters used for MSI data processing in MZmine(v4.2.6)

| **Module Name** | **Key Parameters** | **Contents** |
| --- | --- | --- |
| Mass detection | MS level | MS1 |
|  | Mass detection | Auto |
|  | Noise level | 1.0*10^3^ |
| Image builder | m/z tolerance | 0.005m/z or 20ppm |
|  | Minimum absolute height | 5.0*10^3^ |
|  | Minimum total signals | 50 |
|  | Minimum consecutive scans | 5 |
|  | Suffix | images |

##### Table S5. The software source of this work.

| **Project Name** | **Home Page** |
| --- | --- |
| AirdPro | https://github.com/CSi-Studio/AirdPro |
| AirdSDK | https://github.com/CSi-Studio/Aird-SDK |
| Aird-MZmine | https://github.com/CSi-Studio/mzmine3 |
